# UT-018 Accelerates Wound Repair and Hair Follicle Regeneration in Murine Models

**DOI:** 10.64898/2026.05.18.726121

**Authors:** Uday Saxena, K Saranya, Priyanka Jadhav, Shreya Shahapur, Sadiya Mehboob, Gopi Kadiyala, Markendeya Gorantla

## Abstract

UT-018, a stem cell chemoattractant formulation, demonstrated significant regenerative activity across independent murine wound-healing and hair-regeneration studies. Topical treatment accelerated wound closure, enhanced granulation tissue formation, improved collagen organization, increased fibroblast proliferation, and enhanced dermal vascularization. Separate hair-growth studies demonstrated increased follicular density, deeper follicular penetration, enhanced dermal vascularization, and induction of anagen-phase transition by UT-018. Mechanistic studies demonstrated strong intracellular cAMP generation and activation-associated β-catenin phosphorylation consistent with GPCR-mediated regenerative signaling.

## Introduction

Cutaneous wound healing and follicular regeneration require coordinated epithelial repair, extracellular matrix remodeling, angiogenesis, fibroblast activation, and stem/progenitor-cell recruitment. These regenerative processes are tightly regulated through interconnected signaling pathways that control cellular migration, proliferation, differentiation, and tissue remodeling following injury. Despite advances in regenerative medicine, currently available therapies for chronic wounds and hair growth remain limited by incomplete regenerative activity, poor restoration of tissue architecture, and inability to effectively activate endogenous regenerative signaling pathways.

β-catenin signaling plays a central role in tissue regeneration, epithelial repair, and hair follicle cycling. In skin biology, β-catenin signaling is associated with wound healing, angiogenesis, extracellular matrix remodeling, and induction of the anagen phase of hair growth. In addition to canonical Wnt signaling, β-catenin activity can also be modulated through GPCR/cAMP/PKA-associated pathways, including phosphorylation at activation-associated sites such as Ser552 and Ser675, supporting regenerative transcriptional responses involved in tissue repair and follicular regeneration.

UT-018, consisting of selected bioactive amino acids combined in a defined ratio, was evaluated as a regenerative signaling composition capable of activating phenotypic and molecular events associated with tissue repair and follicular regeneration. The present studies evaluated the effects of UT-018 in independent murine wound-healing and hair-regeneration models together with mechanistic studies examining intracellular cAMP generation and activation-associated phospho-β-catenin signaling.

## Materials and Methods

### Full-Thickness Excisional Wound Model

6-mm dorsal full-thickness wounds were created under isoflurane anesthesia using a sterile biopsy punch. UT-018 (25 mM, 50 µL/wound) was applied topically twice daily for 14 days; controls received PBS base only. Endpoints included wound closure (digital photographs analyzed in ImageJ), histology (H&E, Masson’s trichrome).

Endpoints included:

- Percentage wound closure
- Residual wound area
- Healing rate
- Histological assessment
- Granulation tissue formation
- Collagen organization
- Inflammation

### Hair-Regeneration Studies

C57BL/6 mice (n = 5/group) were depilated to synchronize follicles into telogen phase prior to treatment initiation. UT-018 (25 mM) was applied topically twice daily for 28 days. Vehicle (PBS)-treated animals served as controls.

Endpoints included visual assessment of hair regrowth, histological analysis of follicular density and depth, dermal vascularization, dermal thickness, and identification of anagen-phase induction.

All animal studies were conducted in accordance with the Institutional Animal Ethics Committee (IAEC) and complied with CPSCEA guidelines.

### cAMP Quantification

Intracellular cAMP levels were quantified using a competitive commercial immunoassay. cAMP concentrations were interpolated from standard calibration curves.

### Western Blot Analysis

β-catenin phosphorylation studies were conducted using phospho-specific antibodies obtained commercially and western blot analysis performed to evaluate activation-associated phospho-β-catenin species corresponding to Ser552 and Ser675 phosphorylation-associated bands.

## Results

### UT-018 Accelerates Closure of Wounds

We first examined whether UT-018 could accelerate wound repair in wound-healing model. Compared with PBS-treated controls, UT-018-treated wounds showed earlier separation from control healing kinetics, with visible divergence beginning by Day 3 and persisting through the study period. The UT-018 group showed faster wound contraction, lower residual wound area, and a stronger healing rate profile. UT-018 significantly accelerated wound closure compared to vehicle. At day 7, closure was ∼55–60% in UT-018 vs 30–35% in vehicle (p < 0.01). At day 10, closure reached ∼85–90% in UT-018 vs ∼65–70% in vehicle (p < 0.05). By day 14, nearly complete closure (>95%) was observed in UT-018, while vehicle wounds remained ∼85% closed (Figure 1).

**Figure 1A–B.**
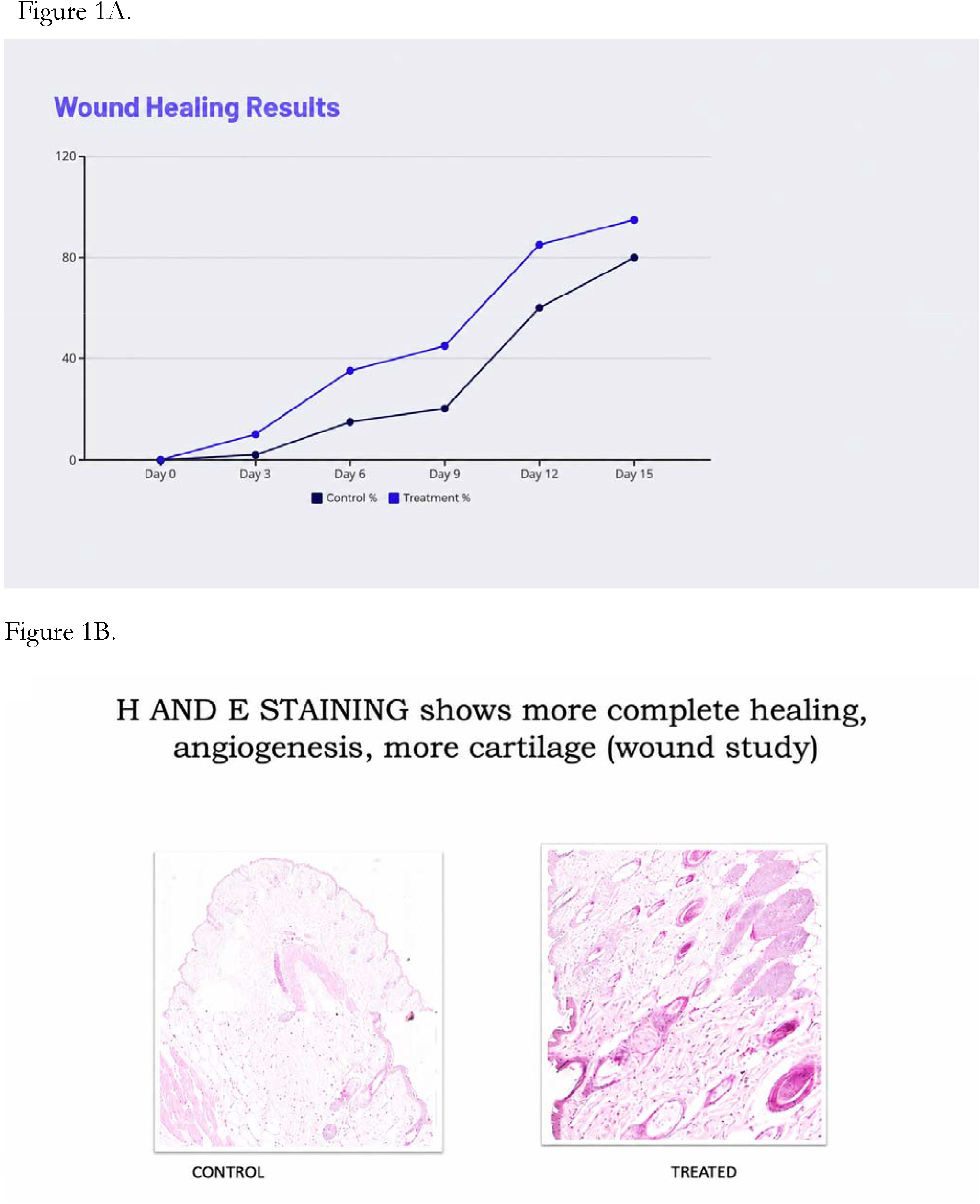
Wound closure kinetics and histological evaluation following topical UT-018 treatment. Wound healing kinetics and representative histological sections shown here demonstrate accelerated wound contraction and improved tissue repair after topical UT-018 treatment. UT-018-treated wounds showed enhanced granulation tissue formation, organized collagen remodeling, increased angiogenesis, and near-complete re-epithelialization compared with PBS-treated controls.

Histological evaluation further showed a more mature repair response, including enhanced granulation tissue formation, more organized collagen remodeling, increased fibroblast proliferation, enhanced vascularization, and near-complete re-epithelialization. Histology also showed earlier re-epithelialization, thicker stratified epidermis, reduced neutrophilic infiltration revealed denser, more mature collagen fibers (Figure 1B.). These findings indicate that UT-018 did not simply reduce wound size, but improved the overall quality and organization of tissue repair. No adverse systemic or local effects were observed.

### Hair-Growth and Follicular-Regeneration Studies

We then examined whether the regenerative effects of UT-018 extended beyond wound closure into follicular regeneration. In depilated C57BL/6 mice, UT-018-treated skin showed earlier visible hair regrowth compared with vehicle-treated controls. Histological analysis demonstrated a clear increase in follicular density, with approximately 52 follicles per field in UT-018-treated sections (PBS treated control animal histopathology showed 40 follicles per field). Follicles were more widely distributed through the dermis and appeared at multiple developmental stages, consistent with activation of follicular cycling and anagen-phase transition. The dermal compartment also showed increased fibroblast proliferation, preserved collagen architecture, minimal inflammation, and enhanced vascularization. These findings suggest that UT-018 promotes follicular regeneration together with organized stromal remodeling without evidence of overt tissue irritation. A summary of these data is shown in Table 1 below and representative histopathology sections are shown in Figure 2 below.

**Table 1.**
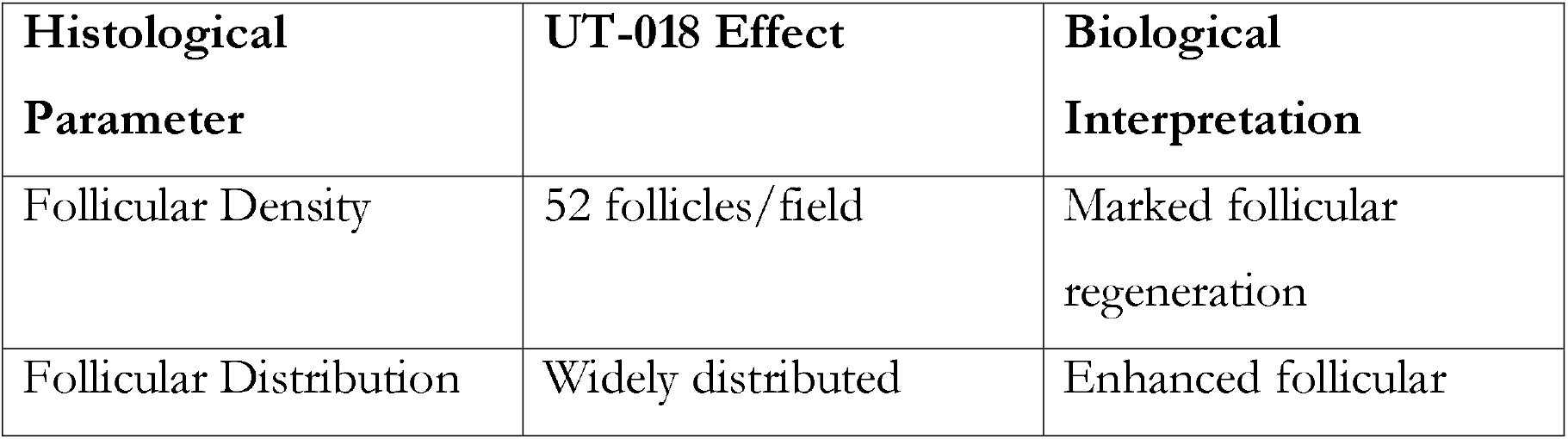
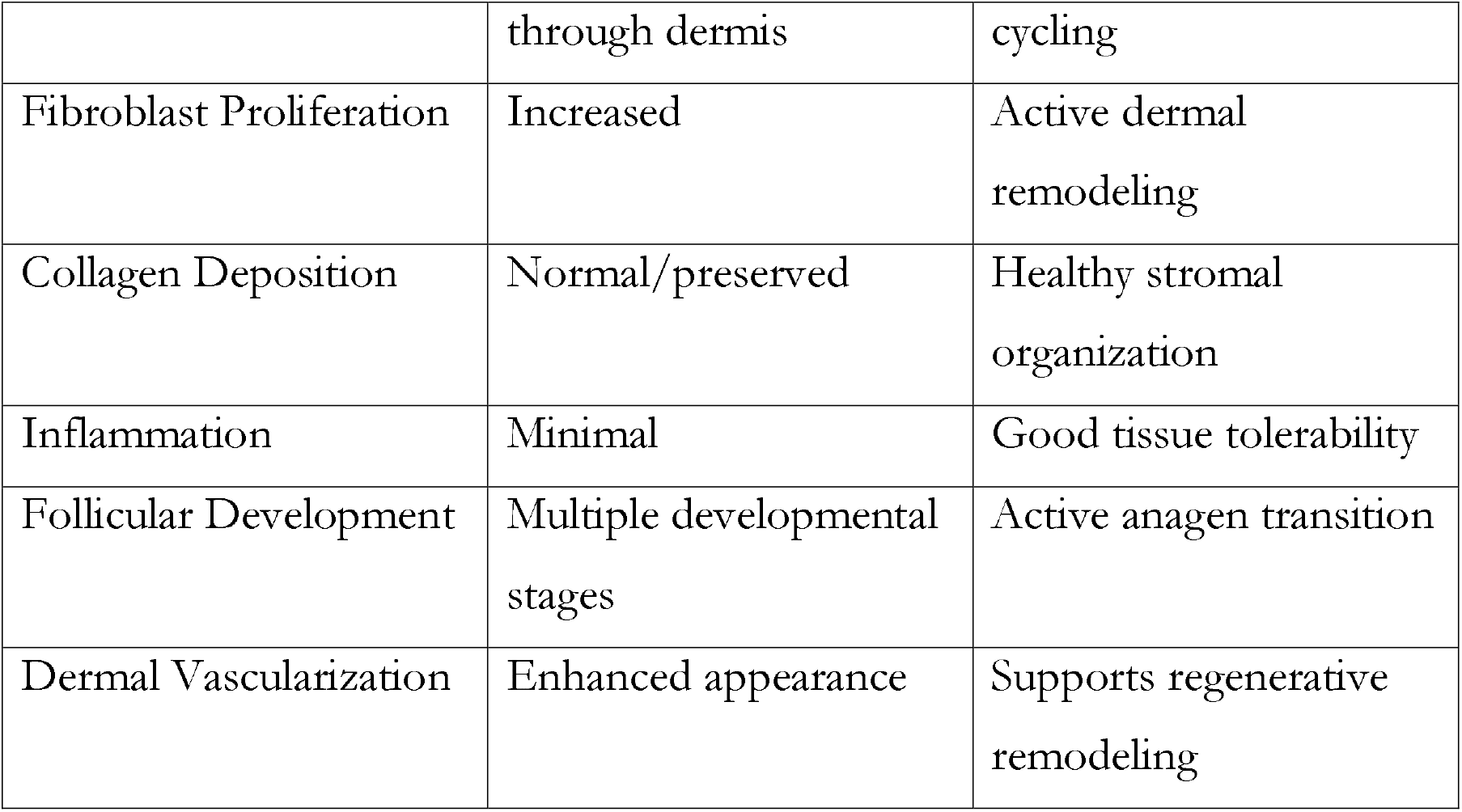

**Figure 2.**
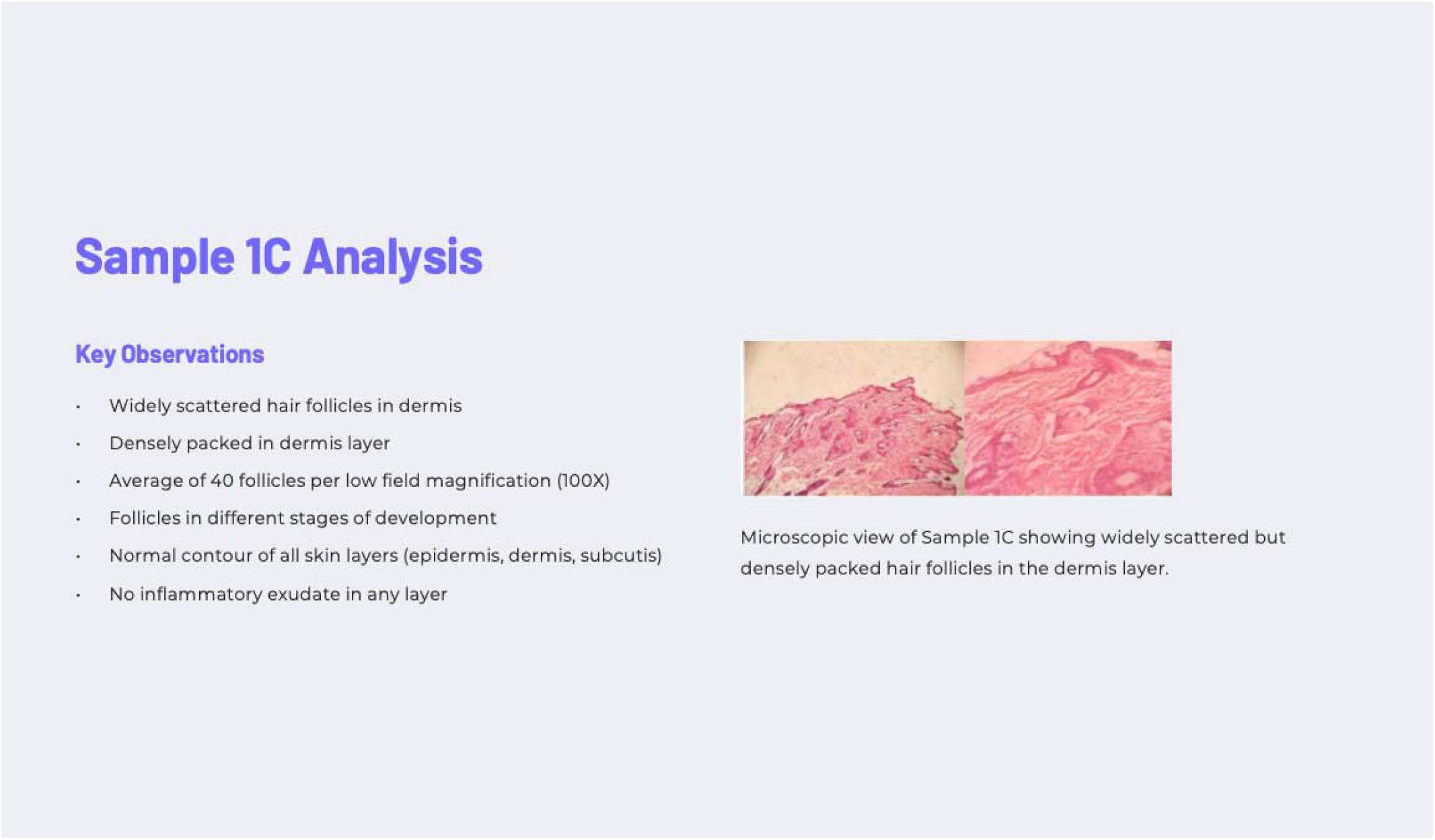
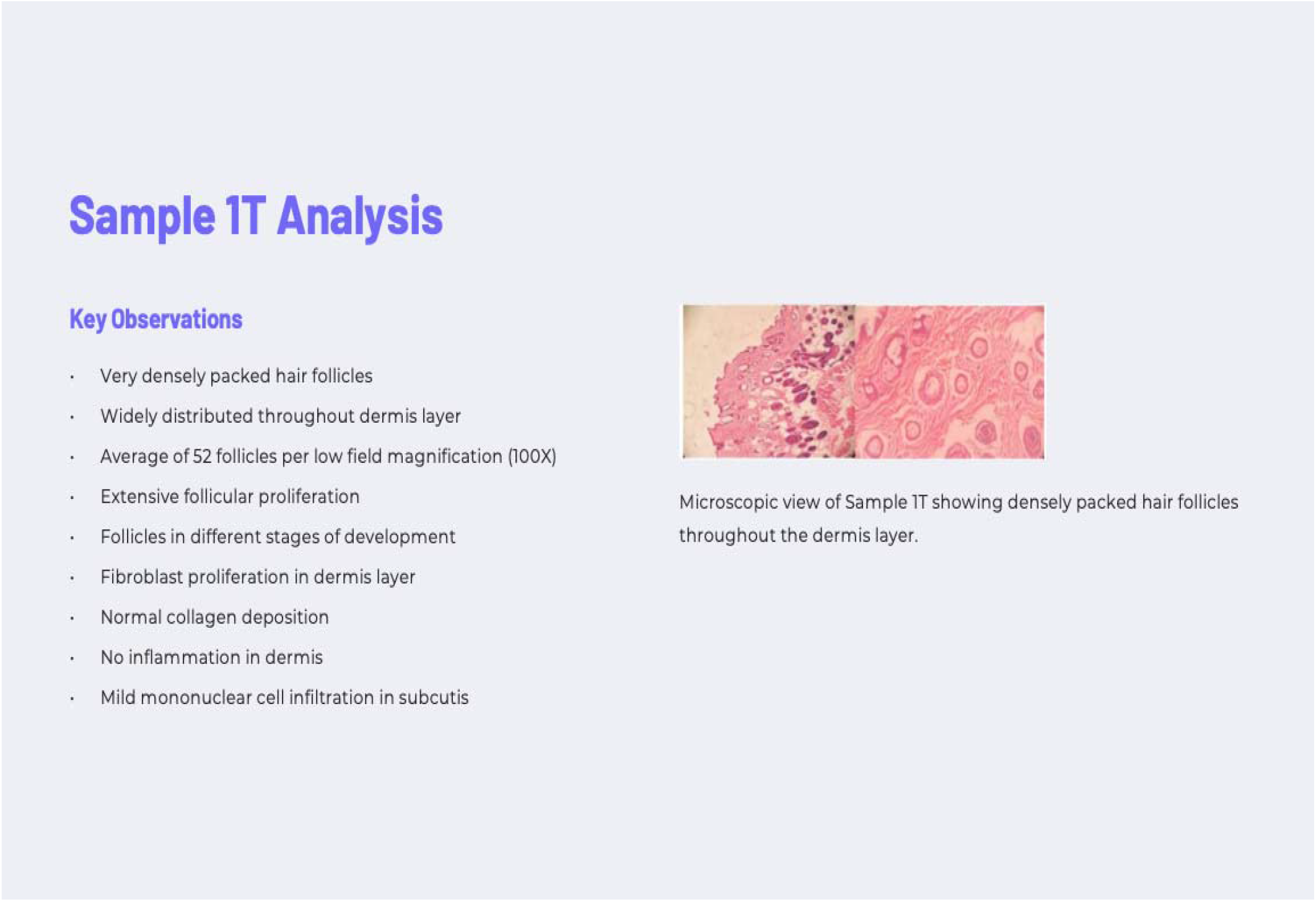
Hair-growth histopathology and follicular-regeneration endpoints. Representative histological sections demonstrate dense follicular packing, deeper follicular penetration into the dermis, increased fibroblast proliferation, preserved collagen architecture, minimal inflammation, and enhanced dermal vascularization following topical UT-018 treatment. Representative control (1 C) and UT-018-treated (1 T) histopathology sections are shown above.

### cAMP increase and β-Catenin Western Blot

We next investigated whether the regenerative phenotypes observed in wound-healing and hair-growth studies were associated with activation of intracellular signaling pathways linked to stem/progenitor-cell recruitment and tissue repair. Intracellular cAMP is a central second messenger involved in regulation of cellular repair, migration, proliferation, and regenerative signaling. Activation of GPCR-mediated cAMP signaling can stimulate protein kinase A (PKA)-dependent pathways, including activation-associated phosphorylation and stabilization of β-catenin, thereby promoting tissue regeneration and follicular repair.

Treatment with UT-018 produced a marked increase in intracellular cAMP levels, demonstrating approximately a **six-fold** elevation relative to untreated controls (Figure 3), consistent with activation of a GPCR-mediated signaling cascade. Elevated cAMP signaling is known to activate protein kinase A (PKA), a central regulator of β-catenin stabilization and regenerative transcriptional programs.

**Figure 3.**
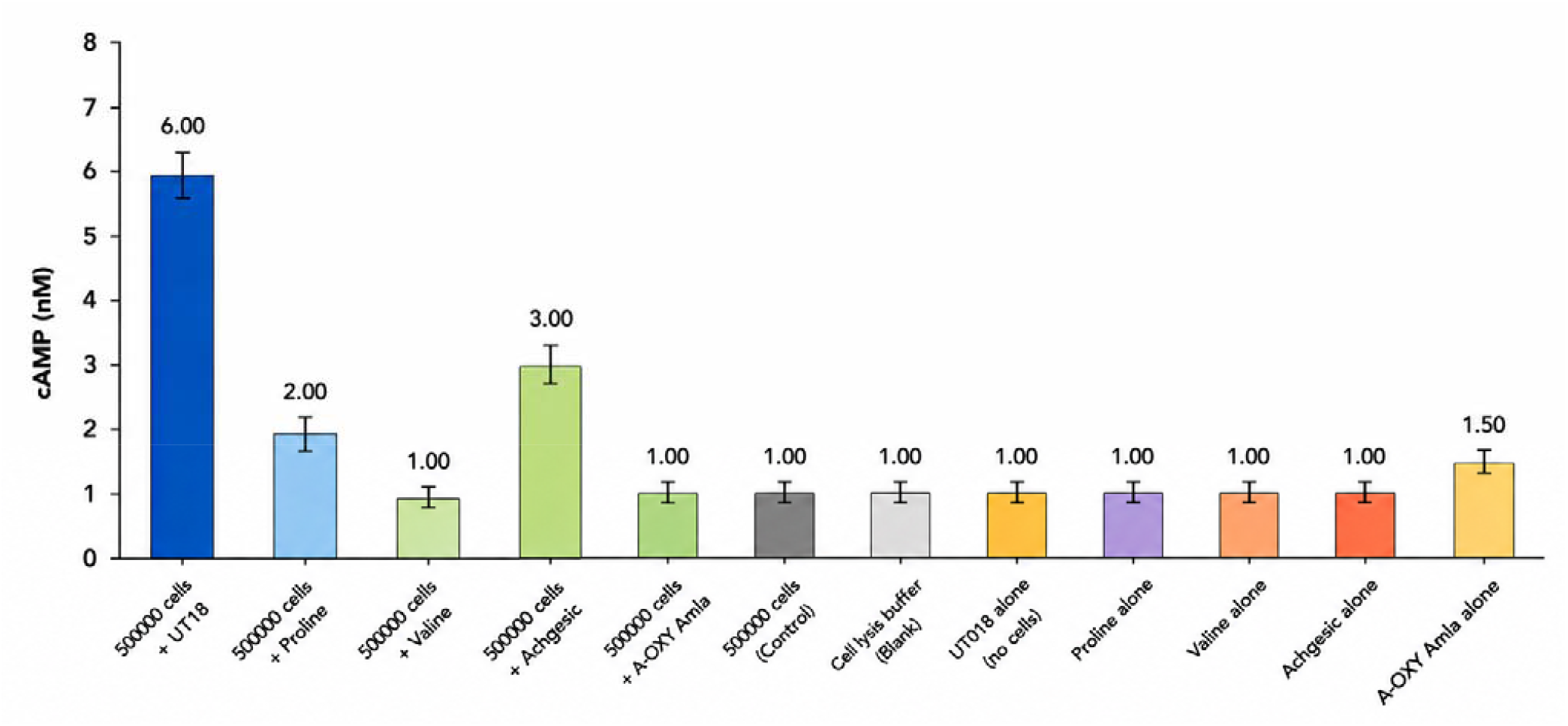
Intracellular cAMP induction following UT18 treatment compared with individual amino acid controls. Cells (5 × 10^5^ per sample) were treated with UT18 or individual amino acid formulations for 30 minutes, followed by intracellular cAMP quantification using a competitive cAMP immunoassay. Absorbance was measured at 450 nm and cAMP concentrations were interpolated from a standard calibration curve. UT18 treatment produced the highest cAMP response (∼6 nM), whereas individual amino acid formulations produced minimal or modest cAMP induction (approximately 1 – 3 nM). UT018 alone (no cells) served as assay control.

β-catenin is a central regulator of tissue regeneration, epithelial repair, and hair follicle cycling. In skin biology, β-catenin signaling is associated with wound healing, angiogenesis, extracellular matrix remodeling, and induction of the anagen phase of hair growth. β-catenin activity can also be modulated through GPCR/cAMP/PKA-associated pathways, including phosphorylation at activation-associated sites such as Ser552 and Ser675, supporting regenerative transcriptional responses.

We therefore explored the effects of UT-018 on phospho-β-catenin species. Western blot analysis demonstrated the appearance of activation-associated phospho-β-catenin species following UT-018 exposure using antibodies to phosphorylated β-catenin at Ser552 and Ser675 phosphorylation sites. Lower-molecular-weight phosphorylated β-catenin bands were observed in treated samples, suggesting activation-associated post-translational modification of β-catenin (Figure 4). The phospho-β-catenin banding pattern observed after UT-018 exposure suggests activation-associated β-catenin modification linked to regenerative signaling activity.

**Figure 4.**
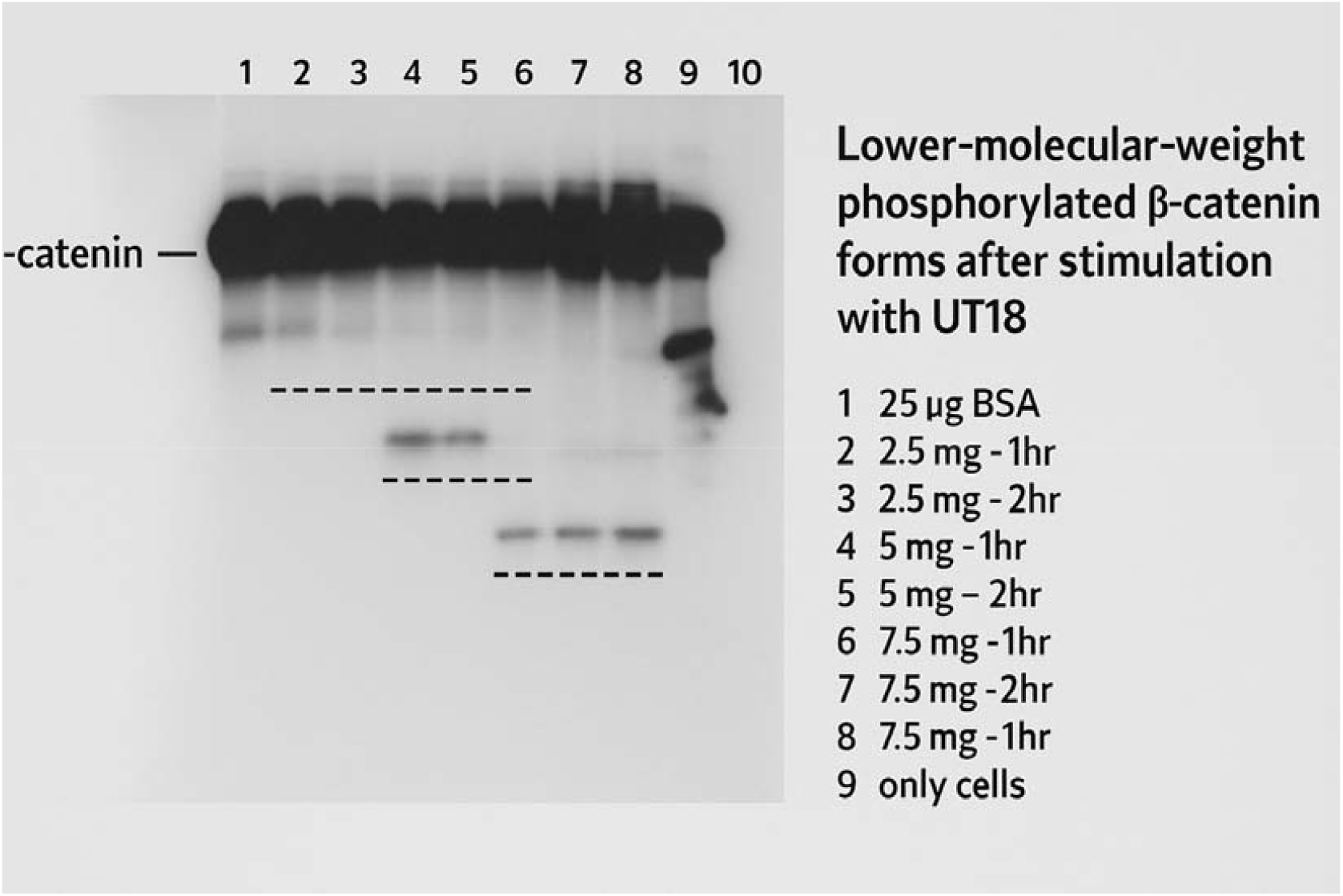
Phospho-β-catenin western blot and regenerative signaling interpretation. Western blot analysis of phosphorylated β-catenin following UT-018 treatment is shown. Increasing amounts of total cellular protein lysate and different incubation time points with UT-018 were loaded across lanes, as indicated in the figure. The upper band corresponds to full-length β-catenin, while lower-molecular-weight phosphorylated β-catenin species appeared selectively after UT-018 stimulation, consistent with activation-associated phosphorylation and/or processing events.

The observed signaling profile was consistent with activation of a GPCR/cAMP-associated pathway linked to β-catenin stabilization and regenerative transcriptional responses.

## Discussion

Across independent wound-healing and follicular-regeneration studies, UT-018 consistently accelerated regenerative repair and improved tissue quality. Histopathological analyses demonstrated enhanced epithelial regeneration, granulation tissue formation, collagen organization, fibroblast proliferation, vascularization, and follicular regeneration.

Mechanistic studies demonstrated approximately six-fold intracellular cAMP elevation and activation-associated β-catenin phosphorylation, supporting activation of GPCR-mediated regenerative signaling pathways. Collectively, the data suggest coordinated activation of pathways associated with angiogenesis, epithelial repair, fibroblast activation, and follicular regeneration.

In conclusion, UT-018 significantly accelerated wound healing and promoted follicular regeneration across independent murine studies. The integrated biological effects included accelerated wound closure, angiogenesis, collagen remodeling, epithelial repair, increased follicular density, and enhanced dermal vascularization. Mechanistic studies suggest involvement of GPCR-mediated cAMP-associated signaling and β-catenin activation supporting a coordinated regenerative signaling mechanism.

## Acknowledgements

We thank the In Vivo biology team at Center for DNA Fingerprinting and Diagnostics (CDFD), an Institute located in Hyderabad, India, for helping us with studies evaluating the effects of UT-018 in murine models. All in vivo studies were conducted in accordance with the Institutional Animal Ethics Committee (IAEC) and complied with CPSCEA guidelines.

